# Tree Diet: Reducing the Treewidth to Unlock FPT Algorithms in RNA Bioinformatics

**DOI:** 10.1101/2021.04.30.442158

**Authors:** Bertrand Marchand, Yann Ponty, Laurent Bulteau

**Affiliations:** LIX CNRS UMR 7161, Ecole Polytechnique, Institut Polytechnique de Paris, Palaiseau, France; LIGM, CNRS, Univ Gustave Eiffel, F77454 Marne-la-Vallée, France

**Keywords:** RNA, treewidth, FPT algorithms, RNA design, structure sequence alignment

## Abstract

Hard graph problems are ubiquitous in Bioinformatics, inspiring the design of specialized Fixed-Parameter Tractable algorithms, many of which rely on a combination of tree-decomposition and dynamic programming. The time/space complexities of such approaches hinge critically on low values for the treewidth *tw* of the input graph. In order to extend their scope of applicability, we introduce the Tree-Diet problem, *i.e*. the removal of a minimal set of edges such that a given tree-decomposition can be slimmed down to a prescribed treewidth *tw*′. Our rationale is that the time gained thanks to a smaller treewidth in a parameterized algorithm compensates the extra post-processing needed to take deleted edges into account.

Our core result is an FPT dynamic programming algorithm for Tree-Diet, using 2^*O*(*tw*)^*n* time and space. We complement this result with parameterized complexity lower-bounds for stronger variants (e.g., NP-hardness when *tw*′ or *tw* − *tw*′ is constant). We propose a prototype implementation for our approach which we apply on difficult instances of selected RNA-based problems: RNA design, sequence-structure alignment, and search of pseudoknotted RNAs in genomes, revealing very encouraging results. This work paves the way for a wider adoption of tree-decomposition-based algorithms in Bioinformatics.

## 1 Introduction

Graph models and parameterized algorithms are found at the core of a sizable proportion of algorithmic methods in bioinformatics addressing a wide array of subfields, spanning sequence processing [1], structural bioinformatics [2], comparative genomics [3], phylogenetics [4], and further examples that can be found in a review by Bulteau and Weller [5]. RNA bioinformatics is no exception, with the prevalence of the secondary structure, an outer planar graph [6], as an abstraction of RNA conformations, and the notable utilization of graph models to represent complex topological motifs called pseudoknots [7], inducing the hardness of several tasks, such as structure prediction [8, 9, 10], structure alignment [11], or structure/sequence alignment [12]. Such motifs are functionally important and conserved, as witnessed by their presence in the consensus structure of 336 RNA families in the 14.5 edition of the RFAM database [13]. Moreover, methods in RNA bioinformatics [14] are increasingly considering non-canonical base pairs and modules [15, 16], further increasing the density of RNA structural graphs and outlining the need for scalable algorithms.

A parameterized complexity approach can be used to circumvent the frequent NP-hardness of relevant problems. It generally considers one or several parameters, whose values are naturally bounded (or much smaller than the input size) within real-life instances. Once relevant parameters have been identified, one aims to design a Fixed Parameter Tractable (FPT) algorithm, having polynomial complexity for any fixed value of the parameter, and reasonable dependency on the parameter value. The treewidth is a classic parameter for FPT algorithms, and intuitively captures a notion of distance of the input to a tree. It is popular in bioinformatics due to the existence of efficient heuristics [17, 18] for computing tree-decompositions of reasonable treewidth. Given a tree-decomposition, many combinatorial optimization tasks can be solved using dynamic programming (DP), in time/space complexities that remain polynomial for any fixed treewidth value. Resulting algorithms remain correct upon (almost) arbitrary modifications of the objective function parameters, and can be adapted to study statistical properties of search spaces through changes of algebra.

Unfortunately, the existence of a parameterized (or FPT) algorithm does not necessarily imply that of a practically-efficient implementation, even when the parameter takes low typical values. Indeed, the dependency of the complexity on the treewidth may be prohibitive, both in terms of time and memory requirements. This limitation is particularly obvious while searching and aligning structured RNAs, giving rise to an algorithmic problem called RNA structure-sequence alignment [19, 20, 12], for which the best known exact algorithm is in Θ(*n.m*^*tw*+1^), with *n* the structure length, *m* the sequence/window length, and *tw* the treewidth of the structure (inc. backbone). Such a complexity becomes impractical for structures having a treewidth higher than *∼* 4, which represent 50 to 70% of known RNA structures, as shown by Figure 1, based on a broad analysis of structures found in the PDB database. Similar complexities hold for problems that can be expressed as (weighted) constraint satisfaction problems, with *m* representing the cardinality of the variable domains. Such frameworks are frequently used for molecular design, both in proteins [21] and RNA [22], and may require the consideration of tree-widths as high as 20 or more [23].

**Figure 1.**
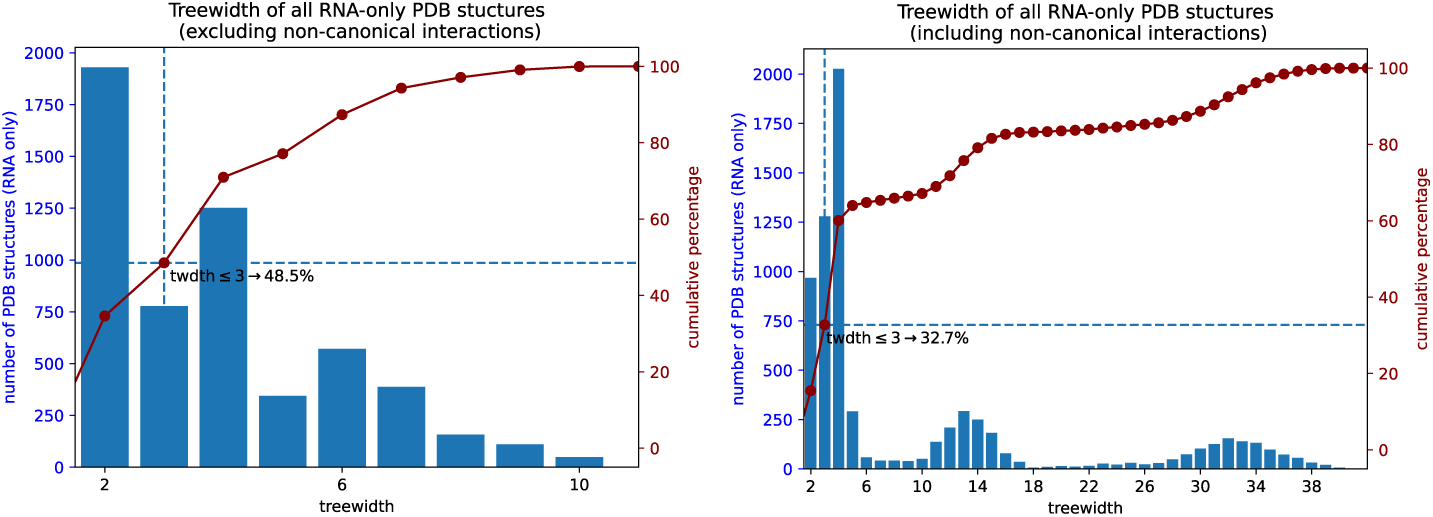
Histogram of treewidth values over all RNA-only structures in the PDB database [24]. The data consists of 5 760 non-redundant graphs, each corresponding to a “chain” of a PDB entity. The nucleotide chains and their base pairs were extracted using the DSSR tool [25]. On each of these graphs, 4 standard treewidth heuristics from the LibTW library [26] (min-degree, min-fill-in, lex-BFS, max-cardinality-search) were launched, and the best width result was selected. Even if these heuristics reputedly tend to yield results close to the optimal, these results are still upper bounds. For each individual structure, the actual treewidth value may be lower. Depending on whether non-canonical base pairs are taken into account (right) or not (left), the proportion of structures having a width *≥* 4 ranges from 50 to 70%. For such values, the complexity of structure-sequence alignment (*O*(*n · m*^*tw*+1^)) becomes prohibitive. It is also worth noting that only *pseudo-knotted* structures may have a treewidth *≥* 3.

In this paper, we investigate a pragmatic strategy to increase the practicality of parameterized algorithms based on the treewidth parameter [27]. We put our instance graphs on a diet, *i.e*. we introduce a preprocessing that reduces their treewidth to a prescribed value by removing a minimal cardinality set of edges. As discussed previously, the practical complexity of many algorithms greatly benefits from the consideration of simplified instances, having lower treewidth. Moreover, specific countermeasures for errors introduced by the simplification can sometimes be used to preserve the correctness of the algorithm. For instance, for searching structured RNAs using RNA structure-sequence alignment [19], an iterated filtering strategy could use instances of increasing treewidth to restrict potential hits, weeding them early so that a – costly – full structure is reserved to (quasi-)hits. This strategy could remain exact while saving substantial time. Alternative countermeasures could be envisioned for other problems, such as a rejection approach to correct a bias introduced by simplified instances in RNA design. An overview of our approach is sketched on Figure 2

**Figure 2.**
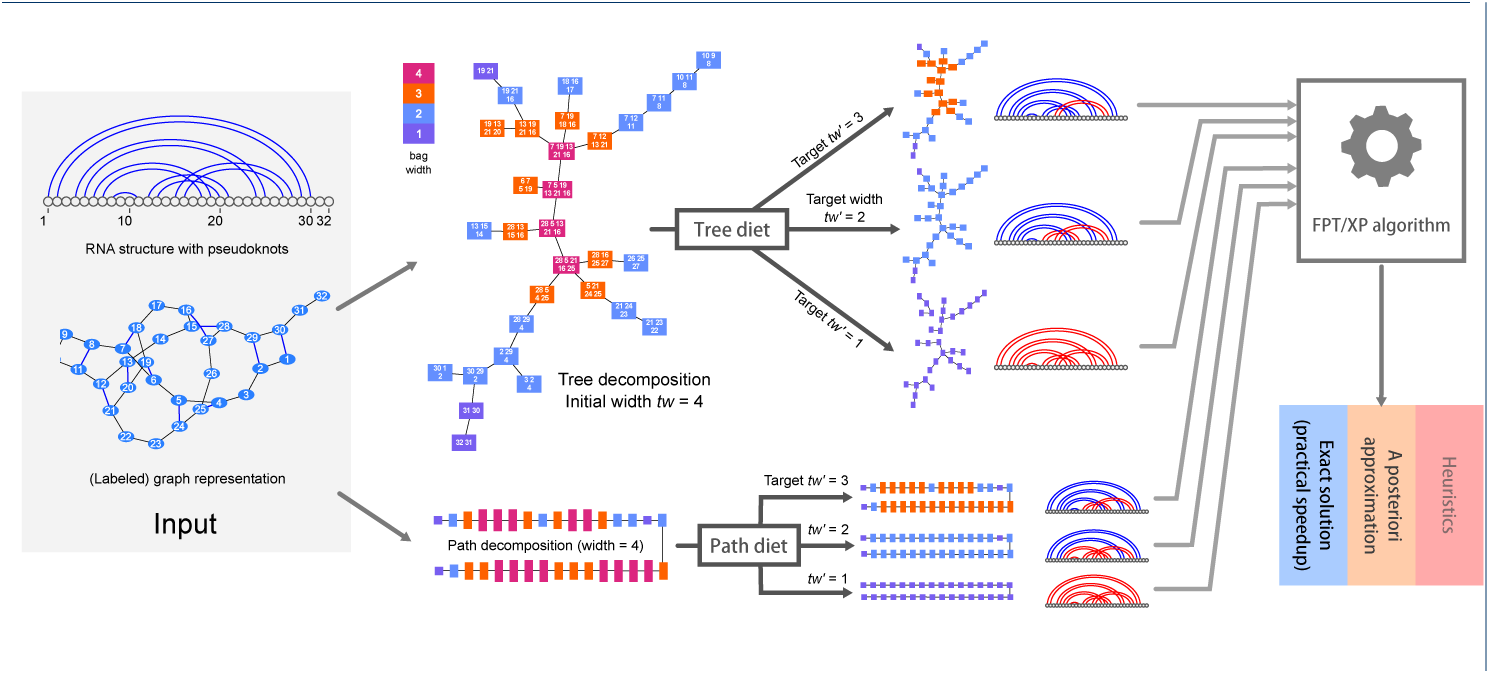
General description of our approach and rationale. Starting from a structured instance, *e.g*. an RNA structure with pseudoknots, our tree-diet/path-diet algorithms extract simplified tree/path decompositions, having prescribed target width *tw*′. Those can be used within existing parameterized algorithms to yield efficient heuristics, *a posteriori* approximations or even exact solutions.

After stating our problem(s) in Section 2, we study in Section 3 the parameterized complexity of the Graph-Diet problem, the removal of edges to reach a prescribed treewidth. We propose, in Section 4, a practical Dynamic Programing FPT algorithm for Tree-Diet, along with possible further optimizations for Path-Diet, two natural simplifications of the Graph-Diet problem, where a tree (resp. path) decomposition is provided as input and used as a guide. Finally, we show in Section 5 how our algorithm can be used to extract hierarchies of graphs/structural models of increasing complexity to provide alternative sampling strategies for RNA design, and speed-up the search for pseudoknotted non-coding RNAs. We conclude in Section 6 with future considerations and open problems.

## 2 Statement of the problem(s) and results

A *tree-decomposition 𝒯* (over a set *V* of vertices) is a tree whose nodes are subsets of *V*, known as bags. The bags containing any *v ∈ V* induce a (connected) subtree of *𝒯*. A *path-decomposition* is a tree-decomposition whose underlying tree *𝒯* is a path. The *width* of *𝒯* (denoted *w*(*𝒯*)) is the size of its largest bag minus 1. An edge *{u, v}* is *visible* in *𝒯* if some bag contains both *u* and *v*, otherwise it is *lost. 𝒯* is a *tree-decomposition of G* if all edges of *G* are visible in *𝒯*. The *treewidth* of a graph *G* is the minimum width over all tree-decompositions of *G*.

**Problem** (Graph-Diet) *Given a graph G* = (*V, E*) *of treewidth tw, and an integer tw*′ *< tw, find a tree-decomposition over V of width at most tw*′ *losing a minimum number of edges from G*.

A *tree-diet* of *𝒯* is any tree-decomposition *𝒯* ′ obtained by removing vertices from the bags of *𝒯*. *𝒯* ′ is a *d*-tree-diet if *w*(*𝒯* ′) *≤ w*(*𝒯*) − *d*.

**Problem** (Tree-Diet) *Given a graph G, a tree-decomposition 𝒯 of G of width tw, and an integer tw*′ *< tw, find a* (*tw* − *tw*′)*-tree-diet of 𝒯 losing a minimum number of edges*.

Note that for Tree-Diet, *𝒯* does not have to be optimal, so the width *tw* of the input tree decomposition might be larger than the actual treewidth of *G*, thus Tree-Diet can be used to reduce the width of *any* input decomposition. We define Binary-Tree-Diet and Path-Diet analogously, where *𝒯* is restricted to be a binary tree (respectively, a path). An example of an instance of Graph-Diet and of Tree-Diet are given in Figure 3.

**Figure 3.**
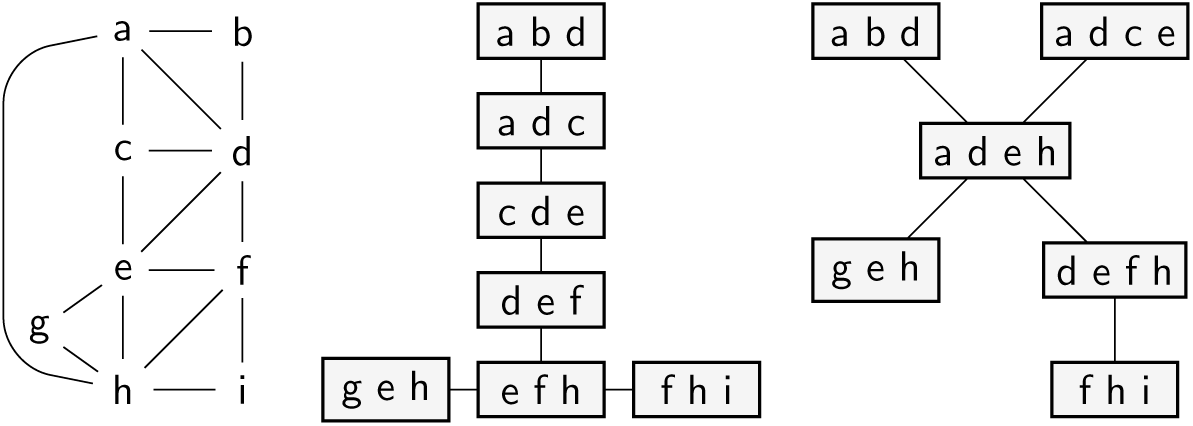
Illustrations for the Graph-Diet and Tree-Diet problems. Given a graph *G* on the left (treewidth 3), an optimal solution for Graph-Diet, with target treewidth 2, yields the tree-decomposition in the middle (edge ah is lost). On the other hand, any 1-tree-diet for the tree-decomposition on the right loses at least 3 edges.

### Parameterized Complexity in a Nutshell

The basics of parameterized complexity can be loosely defined as follows (see [28] for the formal background). A *parameter k* for a problem is an integer associated with each instance which is expected to remain small in practical instances (especially when compared to the input size *n*). An exact algorithm, or the problem it solves, is FPT if it takes time *f* (*k*)poly(*n*), and XP if it takes time *n*^*g*(*k*)^ (for some functions *f, g*). Under commonly accepted conjectures (see for instance [29] for details), W[1]-hard problems may not be FPT, and Para-NP-hard problems (NP-hard even for some fixed value of *k*) are not FPT nor XP.

### 2.1 Our results

Our results are summarized in Table 1. Although the Graph-Diet problem would give the most interesting tree-decompositions in theory, it seems unlikely to admit efficient algorithms in practice (see Section 3).

**Table 1.** Parameterized results for our problems. Algorithm complexities are given up to polynomial time factors (*O*^*^ notation), Δ denotes the maximum number of children in the input tree-decomposition. (*) see Theorem 2 statement for a more precise formulation.

| Parameter Problem | Source treewidth<br>$tw$ | | Target treewidth<br>$tw'$ | Difference<br>$d = tw - tw'$ | | |
| --- | --- | --- | --- | --- | --- | --- |
| GRAPH-DIET | FPT<br>via MSO<br>Theorem 1 | | Para-NP-hard<br>$tw' = 2$<br>EDP($K_4$) [30] | Para-NP-hard*<br>$d = 1$<br>Theorem 2 | | |
| TREE-DIET | XP<br>$O^*((6\Delta)^{tw})$<br>Theorem 7 | FPT open | Para-NP-hard<br>$tw' = 1$<br>Theorem 3 | Para-NP-hard<br>$d = 1$<br>Theorem 4 | | |
| BINARY-TREE-DIET | FPT<br>$O^*(12^{tw})$<br>Theorem 7 | | | W[1]-hard<br>Theorem 5 | XP open | |
| PATH-DIET | | | | | XP<br>$O^*(tw^d)$<br>Theorem 8 | |

Thus we focus on the Tree-Diet relaxation, where an input tree-decomposition is given, which we use as a guide/restriction towards a thinner tree-decomposition. Seen as an additional constraint, it makes the problem harder (the case *tw*′ = 1 becomes NP-hard, Theorem 3, although for Graph-Diet it corresponds to the Spanning Tree problem and is polynomial). With parameter *tw* however, it does help reduce the search space. In Theorem 7 we give an *O*((6Δ)^*tw*^Δ^2^*n*) Dynamic Programming algorithm, where Δ is the maximum number of children of any bag in the tree-decomposition. This algorithm can thus be seen as XP in general, but FPT on bounded-degree tree-decompositions (e.g. binary trees and paths). This is not a strong restriction, since the input tree may safely and efficiently be transformed into a binary one (see Supplementary Section A for more details). Moreover, the duplications of bags which are used in the conversion may only decease the number of lost edges incurred by Tree-Diet.

We also consider the case where the treewidth needs to be reduced by *d* = 1 only, this without constraining the source treewidth. We give a polynomial-time algorithm for Path-Diet in this setting (Theorem 8) which generalizes into an XP algorithm for larger values of *d*, noting that an FPT algorithm for *d* is out of reach by Theorem 5. We also show that the problem is Para-NP-hard if the tree degree is unbounded (Theorem 4).

## 3 Algorithmic Limits: Parameterized Complexity Considerations

Graph-Diet can be seen as a special case of the Edge Deletion Problem (EDP) for the family of graphs *ℋ* of treewidth *tw*′ or less: given a graph *G*, remove as few edges as possible to obtain a graph in *ℋ*. Such edge modification problems are more often parameterized by the number *k* of edited edges (see [31] for a complete survey). Given our focus on increasing the practicality of treewdith-based algorithms in bioinformatics, we restrict our focus to treewidth related parameters *tw, tw*′ and *d* = *tw* − *tw*′.

Considering the target treewidth *tw*′, we note that EDP is NP-hard when *ℋ* is the family of treewidth-2 graphs [30], namely *K*_4_-free graphs, hence the notation EDP(*K*_4_). It follows that Graph-Diet is Para-NP-hard for the target treewidth parameter *tw*′.

### 3.1 Graph-Diet: practical solutions seem unlikely

For a combination of the parameters *tw*′ and *k*, we could imagine graph minor theorems yielding parameterized algorithms “for free”, as it is often the case with treewidth-based problems. In this respect, Graph-Diet corresponds to deciding if a graph *G* belongs to the family of graphs having treewidth *tw*′, augmented by *k* additional edges, denoted as Treewidth-*tw*′+*k*e since its introduction by Cai [32]. If this family were minor-closed, polynomial minor-free-testing [33, 34] would yield an FPT algorithm. However, this is not the case: for some graphs in the family, an edge contraction yields a graph *G*′ not in Treewidth-*tw*′+*k*e, as illustrated by Figure 4.

**Figure 4.**
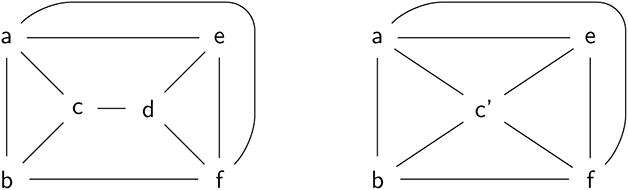
A graph *G* (left) with treewidth 3. Deleting edge cd gives treewidth 2, implying that *G ∈* Treewidth2 + 1e. However, if one contracts edge cd, then the resulting graph (right) has treewidth 3, and deleting any single edge does not decrease the treewidth. This example shows that the graph family Treewidth 2+1e is not minor-closed.

Regarding the source graph treewidth *tw*, the vertex deletion equivalent of Graph-Diet, where one asks for a minimum subset of vertices to remove to obtain a given treewidth, is known as a Treewidth Modulator. This problem has been better-studied than its edge-deletion counterpart [35], and has been shown to be FPT for the treewidth [36]. For the edge-deletion version (Graph-Diet), we can use an optimization variant of Courcelle’s Theorem [29, Thm. 7.12] to show that the problem is FPT for *tw*. However, this is a purely theoretical result as the running-time of such “black-box” algorithms typically involve towers of exponentials on the treewidth parameter.

#### Theorem 1

Graph Diet *is* FPT *for the treewidth*.

*Proof* We formulate Graph Diet as a Monadic Second-Order Logic (MSO) forumula as follows: given a graph *G* = (*V, E*), an integer *tw*′ and a set *X* of edges, let *ϕ*_*tw*_ (*G, X*) be true iff *G*[*E \ X*] has treewidth *tw*′. Clearly *ϕ*_*tw*_ can be expressed as an MSO formula, since both *G*[*E \ X*] and “being of treewidth *tw*′” can be expressed in MSO [37]. Thus, by Arnborg et al. [38], there exists an algorithm that, given *G* of treewidth *tw*, finds a set *X* of minimum size satisfying *ϕ*_*tw*_′ (*G, X*) in time *f*_*tw*_ ′ (*tw*) *· n*. Writing *g*(*tw*) = max_*tw′ ≤tw*_ *f*_*tw*_′ (*tw*), this yields an algorithm for Graph Diet running in time at most *g*(*tw*) *· n*. □

Overall, even though Graph Diet is FPT for the treewidth, “practical” exact algorithms seem out of reach. Indeed, any algorithm for Graph-Diet can be used to compute the Treewidth of an arbitrary graph, for which current state-of-the-art exact algorithms require time in 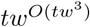 [27]. We thus have the following conjecture, which motivates the Tree-Diet relaxation of the problem.

#### Conjecture 1

Graph-Diet *does not admit algorithms with single-exponential running time for the treewidth*.

On a related note, it is worth noting that Edge Deletion to other graph classes (interval, permutation, …) does admit efficient algorithms when parameterized by the treewidth alone [39], painting a contrasted picture.

Finally, for parameter *d*, any polynomial-time algorithm for constant *d* would allow to compute the treewidth of any graph in polynomial time. Since treewidth is NP-hard we have the following result.

#### Theorem 2

*There is no* XP *algorithm for* Graph-Diet *with parameter d unless* P*=* NP.

*Proof* We consider the decision version of Graph-Diet where a bound *k* on the number of deleted edges is given. We build a Turing reduction from Treewidth: more precisely, assuming an oracle for Graph-Diet with *d* = 1 is available, we build a polynomial-time algorithm to compute the treewidth of a graph *G*. This is achieved by computing Graph-Diet(*G, tw, d* = 1, *k* = 0) for decreasing values of *tw* (starting with *tw* = |*V* |): the first value of *tw* for which this call returns no solution is the treewidth of *G*. Note that this is not a *many-one* reduction, since several calls to Graph-Diet may be necessary (so this does not precisely qualify as an NP-hardness reduction, even though a polynomial-time algorithm for Graph-Diet(*G, tw, d* = 1, *k* = 0) would imply P=NP). □

### 3.2 Lower Bounds for Tree-Diet

Parameters *tw*′ and *d* would be the most interesting in practice, since parameterized algorithms would be efficient for small diets *or* small target treewidth. However, we prove strong lower-bounds for Tree-Diet on each of these parameters, leaving very little hope for parameterized algorithms (we thus narrow down the possible algorithms to the combined parameter *tw*′ + *d*, i.e. *tw*, see Section 4). Only XP for parameter *d* when *𝒯* has a constant degree remains open (cf. Table 1).

#### Theorem 3

Tree-Diet *and* Path-Diet *are* Para-NP*-hard for the target treewidth parameter tw*′ (NP*-hard for tw*′ = 1).

*Proof* By reduction from the NP-hard problem Spanning Caterpillar Tree [40]: given a graph *G*, does *G* have a spanning tree *C* that is a caterpillar? Given *G* = (*V, E*) with *n* = |*V* |, we build a tree-decomposition *𝒯* of *G* consisting of *n* − 1 bags containing all vertices (the width of the decomposition is therefore *n* − 1) connected in a path. Then (*G, 𝒯*) admits a tree-diet to treewidth 1 with *n*−1 visible edges if, and only if, *G* admits a caterpillar spanning tree. Indeed, the subgraph of *G* with visible edges must be a graph with pathwidth 1, i.e. a caterpillar [41]. With *n*−1 visible edges, the caterpillar connects all *n* vertices together, i.e. it is a spanning tree. □

#### Theorem 4

Tree-Diet*is* Para-NP*-hard for parameter d. More precisely, it is* W[1]*-hard for parameter* Δ, *the degree of 𝒯, even when d* = 1.

*Proof* By reduction from Multi-Colored Clique. Consider a *k*-partite graph *G* = (*V, E*) with 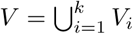. We assume that *G* is regular (each vertex has degree *δ* and that each *V*_*i*_ has the same size *n* (Multi Colored Clique is W[1]-hard under these restrictions [28, 29]). Let 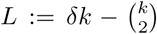 and *N* = max*{*|*V* |, *L* + 1*}*. We now build a graph *G*′ and a tree-decomposition *𝒯* ′: start with *G*′ := *G*. Add *k* independent cliques *K*_1_, …, *K*_*k*_ of size *N* + 1. Add *k* sets of *N* vertices *Z*_*i*_ (*i ∈* [*k*]) and, for each *i ∈* [*k*], add edges between each *v ∈ V*_*i*_ and each *z ∈ Z*_*i*_. Build *𝒯* using 2*k* + 1 bags *T*_0_, *T*_1,*i*_, *T*_2,*i*_ for *i ∈* [*k*], such that *T*_0_ = *V, T*_1,*i*_ = *V*_*i*_ *∪ K*_*i*_ and *T*_2,*i*_ = *V*_*i*_ *∪ Z*_*i*_. The tree-decomposition is completed by connecting *T*_2,*i*_ to *T*_1,*i*_ and *T*_1,*i*_ to *T*_0_ for each *i ∈* [*k*]. Thus, *𝒯* is a tree-decomposition of *G*′ with Δ = *k* and maximum bag size *n* + *N* + 1 (vertices of *V* induce a size-3 path in *𝒯*, other vertices appear in a single bag, edges of *G* appear in *T*_0_, edges of *K*_*i*_ in *T*_1,*i*_, and finally edges between *V*_*i*_ and *Z*_*i*_ appear in *T*_2,*i*_). The following claim completes the reduction:

*𝒯* has a 1-tree-diet losing at most *L* edges from *G*′ *⇔ G* has a *k*-clique.

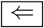Assume *G* has a *k*-clique *X* = *{x*_1_, …, *x*_*k*_*}* (with *x*_*i*_ *∈ V*_*i*_). Build *𝒯* ′ by removing each *x*_*i*_ from bags *T*_0_ and *T*_1,*i*_. Then *𝒯* ′ is a 1-tree-diet of *𝒯*. There are no edges lost by removing *x*_*i*_ from *T*_1,*i*_ (since *x*_*i*_ is not connected to *K*_*i*_), and the edges lost in *T*_0_ are all edges of *G* adjacent to any *x*_*i*_. Since *X* forms a clique and each *x*_*i*_ has degree *δ*, there are 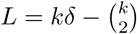 such edges.

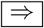Consider a 1-tree-diet *𝒯* ′ of *𝒯* losing *L* edges. Since each bag *T*_1,*i*_ has maximum size, *𝒯* ′ must remove at least one vertex *x*_*i*_ in each *T*_1,*i*_. Note that *x*_*i*_ *∈ V*_*i*_ (since removing *x*_*i*_ *∈ K*_*i*_ would loose at least *N ≥ L*+1 edges). Furthermore, *x*_*i*_ may not be removed from *T*_2,*i*_ (otherwise *N* edges between *x*_*i*_ and *Z*_*i*_ would be lost), so *x*_*i*_ must also be removed from *T*_0_. Let *K* be the number of edges in *G*[*{x*_1_ … *x*_*k*_*}*]. The total number of lost edges in *T*_0_ is *δk* − *K*. Thus, we have *δk* − *K ≤ L* and 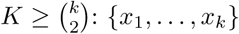 form a *k*-clique of *G*. □

#### Theorem 5

Path-Diet *is* W[1]*-hard for parameter d*.

*Proof* By reduction from Clique. Given a *δ*-regular graph *G* with *n* vertices and *m* edges and an integer *k*, consider the trivial tree-decomposition *𝒯* of *G* with a single bag containing all vertices of *G* (it has width *n* − 1). Then (*𝒯, G*) has a *k*-tree-diet losing 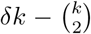 edges if and only if *G* has a *k*-clique. Indeed, such a tree-diet *𝒯*′ would remove a set *X* of *k* vertices from *G* and losing 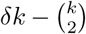 edges, so *X* induces 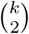 edges and is a *k*-clique of *G*. □

## 4 FPT Algorithm

### 4.1 For general tree-decompositions

We describe here a *O*(3^*tw*^*n*)-space, *O*(Δ^*tw*+2^ *·* 6^*tw*^*n*)-time dynamic programming algorithm for the Tree-Diet problem, with Δ and *tw* being respectively the maximum number of children of a bag in the input tree-decomposition and its width. On *binary* tree-decompositions (where each bag has at most 2 children), it yields a *O*(3^*tw*^*n*)-space *O*(12^*tw*^*n*)-time FPT algorithm.

#### 4.1.1 Coloring formulation

We aim at solving the following problem: given a tree-decomposition *𝒯* of width *tw* of a graph *G*, we want to remove vertices from the bags of *𝒯* to reach a target width *tw*′ while *losing* as few edges from *G* as possible. We tackle the problem through an equivalent *coloring* formulation: our algorithm will assign a color to each occurrence of a vertex in the tree decomposition. We work with three colors: red (r), orange (o), and green (g). Green means that the vertex is kept in the bag, while orange and red means removal of the vertex. An edge is thus visible within a bag when both its ends are green. It is lost if there is no bag where it is visible. To ensure equivalence with the original problem, the colors will be assigned following local rules, which we now describe.

##### Definition 1

*A coloring of vertices in the bags of the decomposition is said to be* valid *if it follows the following rules:*

- *A vertex of a bag not present in its parent may be green or orange (R1)*
- *A green vertex in a bag may be either green or red in its children (R2)*
- *A red vertex in a bag must stay red in its children (R3)*
- *An orange vertex in a bag has to be either orange or green in exactly one child (unless there is no child with this vertex), and must be red in the other children (R4)*

##### These rules are summarized in Figure 6 (a)

When going down the tree, a green vertex may only stay green or permanently become red. An immediate consequence of these rules is therefore that the green occurences of a given vertex form a (possibly empty) connected subtree. Informally, orange vertices are locally absent but “may potentially be found further down the tree”, while red vertices are removed from both the current bag and its entire subtree. Figure 6 (b) shows an example sketch for a valid coloring of the occurrences of a given vertex in the tree-decomposition. A vertex may only be orange along a path starting form its highest occurrence in the tree, with any part branching off that path entirely red. It ends at the top of a (potentially empty) green subtree, whose vertices may also be parents to entirely red subtrees.

**Figure 5.**
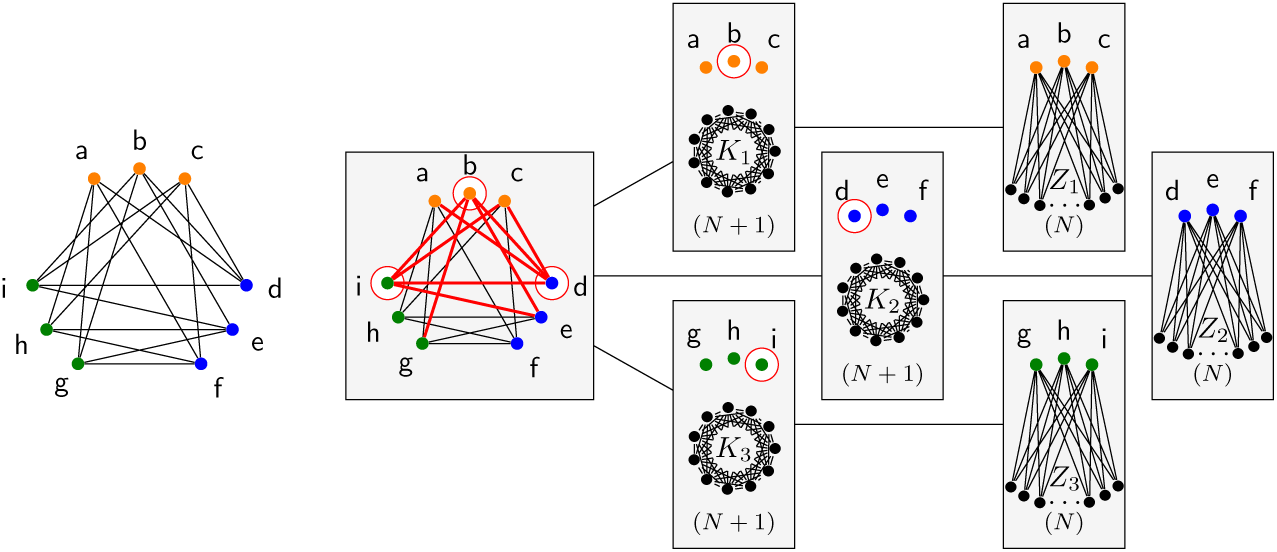
Reduction for Theorem 4 showing that Tree-Diet is NP-hard even for *d* = 1, from a graph *G* (left) with *k* = 3 and *n* = 3 to a graph *G*′ (right, given by its tree-decomposition of width *N* + *n* + 1): a 1-tree-diet for *G*′ amounts to selecting a *k*-clique in the root bag, i.e. in *G*.

**Figure 6.**
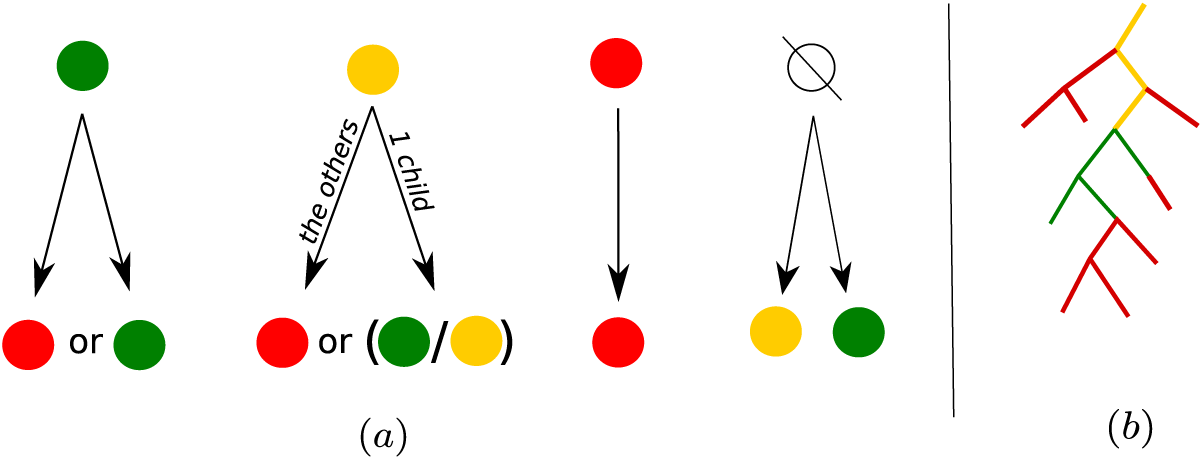
(*a*) Color assignation rules for vertices, when going down-tree. (*b*) Sketch of the general pattern our color assignation rules create on *𝒯*_*u*_, the subtree of bags containing a given vertex *u*

We will now more formally prove the equivalence of the coloring formulation to the original problem. Let us first introduce two definitions. Given a valid coloring *𝒞* of a tree-decomposition of *G*, an edge (*u, v*) of *G* is said to be *realizable* if there exists a bag in which both *u* and *v* are green per *𝒞*. Given an integer *d*, a coloring *𝒞* of *𝒯* is said to be *d*−*diet-valid* if removing red/orange vertices reduces the width of *𝒯* from *w*(*T*) to *w*(*T*) − *d*.

###### Proposition 1

*Given a graph G, a tree-decomposition 𝒯 of width tw, and a target width tw*′ *< tw, The* Tree-Diet *problem is equivalent to finding a* (*tw* − *tw*′)*-dietvalid coloring 𝒞 of 𝒯 allowing for a number of realizable edges in G as large as possible*.

*Proof* Given a (*tw* − *tw*′)-tree-diet of *𝒯*, specifying which vertices are removed from which bags, we obtain a valid coloring *𝒞* for *𝒯* incurring the same number of lost (unrealizable) edges. To start with, a vertex *u* is colored green in the bags where it is not removed. By the validity of *𝒯* ′ as a decomposition, this set of bags forms a connected subtree, that we denote 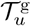. We also write *𝒯*_*u*_ for the subtree of bags containing *u* in the original decomposition *𝒯*. If 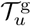 and *𝒯*_*u*_ do not have the same root, then *u* is colored orange on the the path in *𝒯* from the root of *𝒯*_*u*_ (included) and the root of 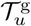 (excluded). Vertex *u* is colored red in any other bag of *𝒯*_*u*_ not covered by these two cases. The resulting coloring follows rules (R1-4) and induces the same set of lost/non-realizable edges as the original (*tw* − *tw*′)-tree-diet. Conversely, an equivalent (*tw* − *tw*′)-tree-diet is obtained from a (*tw* − *tw*′)-diet-valid coloring by removing red/orange vertices and keeping green ones. If a given vertex has no green occurences, it is entirely removed from the tree decomposition and all its edges are *lost* (it becomes an isolated vertex). We may add it back to the tree decomposition by introducing a new bag containing only this vertex, which we connect arbitrarily to the tree decomposition. □

#### 4.1.2 Decomposition of the search space and sub-problems

Based on this coloring formulation, we now describe a dynamic programming scheme for the Tree-Diet problem. We work with sub-problems indexed by tuples (*X*_*i*_, *f*), with *X*_*i*_ a bag of the input tree decomposition and *f* a coloring of the vertices of *X*_*i*_ in green, orange or red (in particular, *f* ^−1^(`g`) denotes the green vertices of *X*_*i*_, and similarly for `o` and `r`).

Let us introduce some notations before giving the definition of our dynamic programming table. Given an edge (*u, v*) of *G*, realizable when coloring a tree-decomposition *𝒯* of *G* with *𝒞*, we write 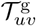 the subtree of *𝒯* in which both *u* and *v* are green. We denote by *𝒯*_*i*_ the subtree of the decomposition rooted at *X*_*i*_, and *C*(*i, f*) the *d*-diet-valid colorings of *𝒯*_*i*_ agreeing with *f* on *i*, with *d* = *tw* − *tw*′. Our dynamic programming table is then defined as:

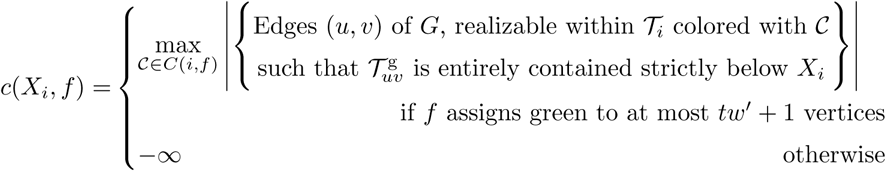

The cell *c*(*X*_*i*_, *f*) therefore aggregates all edges realizable *strictly below X*_*i*_. As we shall see through the recurrence relation below and its proof, edges with both ends green in *X*_*i*_ will be accounted for *above X*_*i*_ in *𝒯*.

We assume w.l.o.g that the tree-decomposition is rooted at an empty bag *R*. Given the definition of the table, the maximum number of realizable edges, compatible with a tree-diet of (*tw* − *tw*′) to *𝒯*, can be found in *c*(*R, ∅*).

The following theorem presents a recurrence relation obeyed by *c*(*X*_*i*_, *f*) :

##### Theorem 6

*For a bag X*_*i*_ *of 𝒯, with children Y*_1_, …*Y*_Δ_, *we have:*

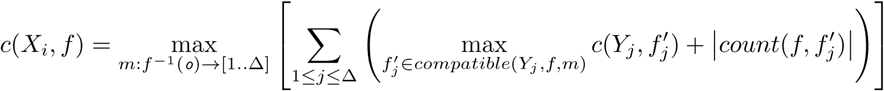

*with*

- *m: a map from the orange vertices in X*_*i*_ *to the children of X*_*i*_. *It decides for each orange vertex u, which child, among those which contain u, will color u orange or green; If there are no orange vertices in X*_*i*_, *only the trivial empty map is considered*.
- *compatible*(*Y*_*j*_, *f, m*): *the set of colorings of Y*_*j*_ *compatible with f on X*_*i*_ *and m;*
- *count*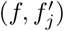: *set of edges of G involving two vertices of Y*_*j*_ *green by* 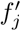, *but such that one of them is either not in X*_*i*_ *or not green by f*.

Note that *compatible*(*Y*_*j*_, *f, m*) may contain colorings 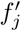 that colour too many vertices in *Y*_*j*_ in green to reach target width *tw*′. In that case 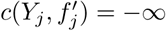.

Theorem 6 relies on the following separation lemma for realizable edges under a valid coloring of a tree-decomposition. Recall that we suppose w.l.o.g that the tree-decomposition is rooted at an empty bag.

##### Lemma 1

*An edge* (*u, v*) *of G, realizable in 𝒯 under 𝒞, is contained in exactly one set of the form count*(*C*_|*P*_, *C*_|*X*_) *with X a bag of 𝒯 and P its parent, C*_|*P*_, *C*_|*X*_ *the restrictions of 𝒞 to P and X, respectively, and “count” defined as above. In addition, X is the root of the subtree of 𝒯 in which both u and v are green*.

*Proof* Given, in a tree-decomposition, a bag *P* colored with *f*, with a child *X* colored with *h*, a more precise definition for *count*(*f, h*) is:

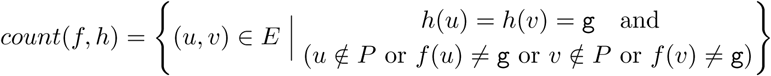

Now, given a realizable edge (*u, v*), in a tree-decomposition *𝒯* colored with *𝒞*, the set of bags in which both *u* and *v* are green forms a connected subtree of *𝒯*. This subtree has a root, or *lowest common ancestor*, that we denote *R*_(*u,v*)_. Since we assumed *𝒯* to be rooted at an empty bag, *R*_(*u,v*)_ is not the root of *𝒯*, and has a parent. We call this parent *P*_(*u,v*)_. Clearly, (*u, v*) belongs to the “count set” associated to the edge (*P*_(*u,v*)_) *→* (*R*_(*u,v*)_) of *𝒯*, while for any other edge *X → Y* of *𝒯*, the colors of *u* and *v* cannot verify the conditions to belong to the associated “count set”. □

##### Proof of Theorem 6

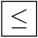 Let us more concisely use *RE*_*↓*_(*𝒯*_*i*_, *𝒞, G*) to denote the set of edges (*u, v*) of *G*, realizable under the (*tw* − *tw*′)-diet-valid coloring *𝒞* of *𝒯*_*i*_, such that 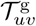 is entirely contained strictly below *X*_*i*_. We have, if *f* contains enough red/orange vertices to reduce the size of *X*_*i*_ to target size:

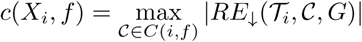 By definition, *c*(*X*_*i*_, *f*) is the maximum number of realizable edges in the subtree-decomposition rooted at *X*_*i*_, such that all green-green occurences of the edge occur strictly below *X*_*i*_, and under the constraint that *f* colors *X*_*i*_. Let *𝒞* be a coloring for *𝒯*_*i*_ realizing the optimum *c*(*X*_*i*_, *f*). Its restrictions to *Y*_1_ … *Y*_Δ_ yield colorings 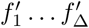. Likewise, its restrictions to the subtree-decompositions 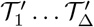 rooted at *Y*_1_ … *Y*_Δ_ yield colorings 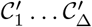 compatible with 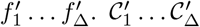 cannot be better than the optimal, so *∀j*, 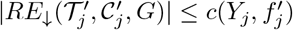 Let (*u, v*) be an edge of *RE*_*↓*_(*𝒯*_*i*_, *𝒞, G*). Per Lemma 1, either (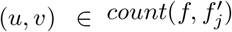 for some *j* (if *Y*_*j*_ is the root of 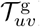) and 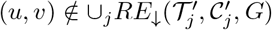 or 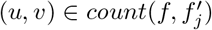 and *∃j* such that 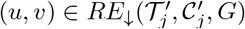. Therefore:

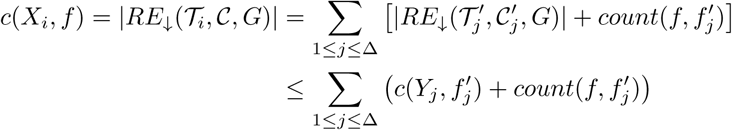

and, a fortiori

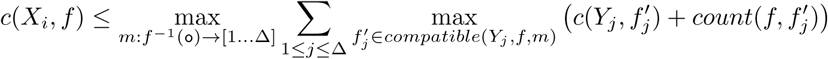
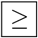 Conversely, given *f*, let *m* be an assignation map for orange vertices and 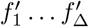 colorings of *Y*_1_ … *Y*_Δ_ compatible with *f* and *m*, and let 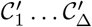 be colotings of 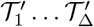 realizing the optima 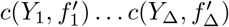. The union of 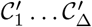 and *f* is a coloring *𝒞* for *𝒯*_*i*_, the subtree-decomposition rooted at *X*_*i*_, which can not be better than optimal (|*RE*_*↓*_(*𝒯*_*i*_, *𝒞, G*)| *≤ c*(*X*_*i*_, *f*)). As before, an edge (*u, v*) either belongs to 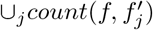 or to 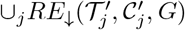 but not both. In any case, it belongs to *RE*_*↓*_(*𝒯*_*i*_, *𝒞, G*). Therefore:

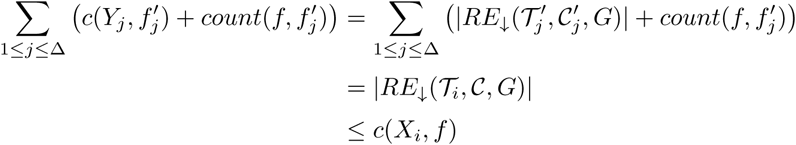

This is true for any choice of *m*, 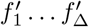, therefore:

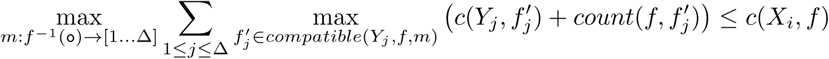

which concludes the proof. □

##### Dynamic programming algorithm

The recurrence relation of Theorem 6 naturally yields a dynamic programming algorithm for the Tree-Diet problem, as stated below:

###### Theorem 7

*There exists a O*(Δ^*tw*+2^ *·* 6^*tw*^ *· n*)*-time, O*(3^*tw*^ *· n*)*-space algorithm for the* Tree-Diet *problem, with* Δ *the maximum number of children of a bag in the input tree-decomposition, and tw its width*.

*Proof of Theorem 7* Given the sub-problems and *c*(*X*_*i*_, *f*)-table definitions, with *R* the (empty) root of the tree-decomposition, *c*(*R, ∅*) is indeed the maximum possible number of realizable edges when imposing a (*tw* − *tw*′)-diet to *𝒯*. The recurrence relation of Theorem 6 therefore lends itself to a dynamic programming approach, over the tree-decomposition *𝒯* following leaf-to-root order, for the problem.

It is reasonable to assume the number of bags in a tree decomposition to be linear in *n* (this is for instance the case for a *nice* tree decomposition [42, 29], or for a tree decomposition obtained from an *elimination ordering*, see [43, 17]). Therefore, the number of entries to the table is *O*(3^*tw*^*n*), given that a bag *X* may be colored in 3^|*X*|^ ways, and that the maximum size of *X* is *tw* + 1. For a given entry *X*_*i*_, one must first enumerate all possible choices of *m* : *f* ^−1^(`o`) *→* [1…Δ], map assigning one child of *X*_*i*_ to each orange vertex in *X*_*i*_. There are *O*(Δ^*tw*+1^) possibilities for *m* in the worst case, as |*f* ^−1^(`o`)| *≤ tw* + 1. Then, for each child *Y*_*j*_, one must enumerate all possible colorings 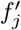 compatible with *f*. Possibilities for 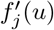 depend on the color by *f* :

- if 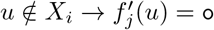 or `g`
- if 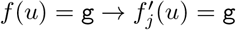 or `r`
- if 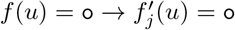 or `g` if 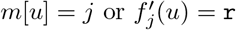 otherwise.
- if 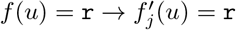

Overall, as there are at most Δ children, *tw* + 1 vertices in each child, and 2 possibilities (see enumeration of cases above) of color for each vertex in a child, yielding a total number of compatible colorings bounded by *O*(Δ *·* 2^*tw*+1^). Multiplying these contributions, the overall time complexity of our algorithm is therefore *O*(Δ^*tw*+2^ *·* 6^*tw*^ *· n*). □

###### Corollary 1

Binary-Tree-Diet (Δ = 2*) admits an* FPT *algorithm for the tw parameter*.

A pseudo-code implementation of the algorithm, using memoization, is included in Supplementary Section B

### 4.2 For path decompositions

In the context of path decompositions, we note that the number of removed vertices per bag can be limited to at most 2*d* without losing the optimality. More precisely, we say that a coloring *𝒞* is *d-simple* if any bag has at most *d* orange and *d* red vertices. We obtain the following result, using transformations given in Figure 7.

**Figure 7.**
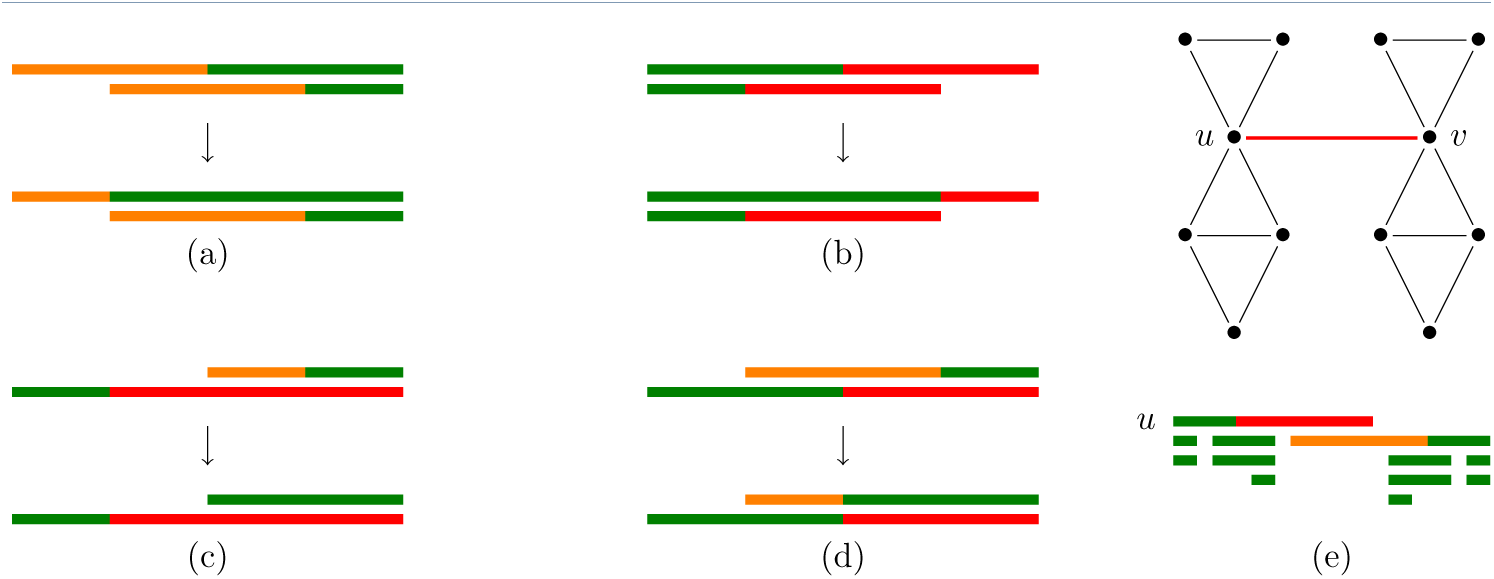
Five cases where two vertices are deleted in the same bag with *d* = 1. Bags are points in the line, and an interval covering all bags containing *v* is drawn for each *v* (with an equivalent coloring, see Proposition 1). Cases (a) to (d) can be safely avoided by applying the given transformations. In the example for case (e), however, it is necessary to delete both vertices u and v form a central bag. It is sufficient to avoid cases (a) and (b) in order to obtain an XP algorithm for *d*.

#### Proposition 2

*Given a graph G and a path-decomposition 𝒯, if 𝒞 is a d-diet-valid coloring of 𝒯 losing k edges, then 𝒯 has a d-diet-valid coloring that is d-simple, and loses at most k edges*.

*Proof of Proposition 2* Consider such a coloring *𝒞* with a maximal number of green vertices. We show that it is *d*-simple. Assume the path-decomposition *𝒯* is rooted in bag *X*_1_ and each *X*_*i*_ is the parent of *X*_*i*+1_. Pick *i* to be the smallest index so that at least *d* + 1 vertices in *X*_*i*_ are colored red by *𝒞*, assume any such *i* exists. Then one of these vertices, say *u*, is not colored red in *X*_*i*−1_ (either because *i* = 1, or it is not in *X*_*i*−1_, or it is orange or green in *X*_*i*−1_). Consider *𝒞*′ obtained by *𝒞* and coloring *u* green in *X*_*i*_. Then *𝒞*′ satisfies local rules R1 through R4 (a green vertex may be absent, green or orange in the parent bag, and a red vertex may be green in the parent bag). Furthermore, it is *d*-diet-valid since it still removes at least *d* (red) vertices in *X*_*i*_. Overall *𝒞*′ is another *d*-diet-valid coloring with more green vertices: a contradiction, so no such *i* exist (and no bag has *d* + 1 red vertices). The same argument works symmetrically for orange vertices. Overall, *𝒞* is *d*-simple. □

Together with Proposition 1, this shows that it is sufficient to restrict our algorithm to *d*-simple colorings. (See also Figure 7). In particular, for any set *X*_*i*_, choosing which *≤ d* vertices are orange and which *≤ d* are red, among the total of *n* vertices, is enough to fix a coloring. The number of such colorings is therefore bounded by *O*(*tw*^2*d*^). Applying this remark to our algorithm presented in Section 4.1 yields the following result:

#### Theorem 8

Path-Diet *can be solved in O*(*tw*^2*d*^*n*)*-space and O*(*tw*^4*d*^*n*)*-time*.

## 5 Proofs of concept

We now illustrate the relevance of our approach, and the practicality of our algorithm for Tree-Diet, by using it in conjunction with FPT algorithms for three problems in RNA bioinformatics. We implemented in C++ the dynamic programming scheme described in Theorem 7 and Supplementary Section B. Its main primitives are made available for Python scripting through pybind11 [44].

It actually allows to solve a generalized *weighted* version of Tree Diet, as explained in Supplementary Section B. This feature allows to favour the conservation of important edges (*e.g*. RNA backbone) during simplification, by assigning them a much larger weight compared to other edges. Our implementation is freely available at https://gitlab.inria.fr/amibio/tree-diet.

The execution time of this implementation on elements of the data set used for Figure 1 (all RNA-only structures of the PDB database) is represented on Figure 8, for input treewidth values of up to 7. It shows that our tree-diet method is applicable with reasonable run-times (≲ 1 hour) for all structures of width *≤* 7. The proofs-of-concepts presented in this section involve however instances with treewidth of up to 9, in the case of RNA design, for which the run-time also stays reasonable.

**Figure 8.**
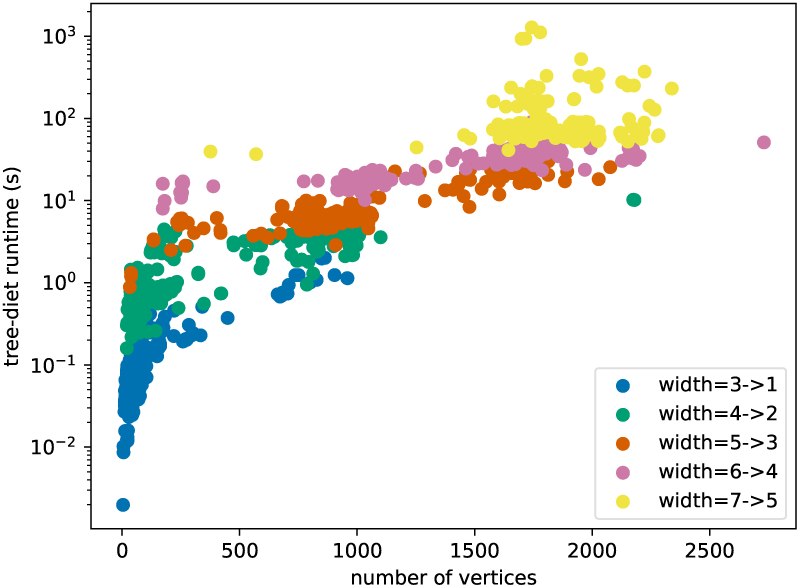
Run-time of the tree-diet algorithm on all RNA-only structures of the PDB database, versus the size (length of the RNA string) of these structures. The data set is the same as Figure 1, limited to structures of treewidth ≤ 7. Structures are colored by their original treewidth. Here, we have asked the algorithm to reduce the treewidth by 2.

### 5.1 Memory-parsimonious unbiased sampling of RNA designs

As a first use case for our simplification algorithm, we strive to ease the sampling phase of a recent method, called RNAPond [22], addressing RNA negative design. The method targets a set of base pairs *S*, representing a secondary structure of length *n*, and infers a set *𝒟* of *m* disruptive base pairs (DBPs) that must be avoided. It relies on a Θ(*k ·* (*n* + *m*)) time algorithm for sampling *k* random sequences (see Supplementary Section C for details) after a preprocessing in Θ(*n · m ·* 4^*tw*^) time and Θ(*n ·* 4^*tw*^) space. Here, the input consists of a graph *G* = ([1, *n*], *S ∪ 𝒟*) and a tree decomposition *𝒯* of *G*, having width *tw*. In practice, the preprocessing largely dominates the overall runtime, even for large values of *k*, and its large memory consumption represents the main bottleneck.

This discrepancy in the complexities/runtimes of the preprocessing and sampling suggests an alternative strategy: relaxing the set of constraints to (*S*′, *𝒟*^*′*^), with (*S*′ *∪ 𝒟*^*′*^) *⊂* (*S ∪ 𝒟*), and compensating it through a rejection of sequences violating constraints in (*S, 𝒟*) *\* (*S*′, *𝒟*^*′*^). The relaxed algorithm would remain unbiased, while the average-case time complexity of the rejection algorithm would be in 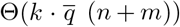 time, where 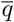 represents the relative increase of the partition function (*≈* the sequence space) induced by the relaxation. The preprocessing step would retain the same complexity, but based on a (reduced) treewidth *tw*′ *≤ tw* for the relaxed graph *G*′ = ([1, *n*], *S*′ *∪ 𝒟*′).

These complexities enable a tradeoff between the rejection (time), and the preprocessing (space), which may be critical to unlock future applications of RNA design. Indeed, the treewidth can be decreased by removing relatively few base pairs, as demonstrated below using our algorithm on pairs inferred for hard design instances.

We considered sets of DBPs inferred by RNAPond over two puzzles in the EteRNA benchmark. The EteRNA22 puzzle is an empty secondary structure spanning 400 nts, for which RNAPond obtains a valid design after inferring 465 DBPs. A tree decomposition of the graph formed by these 465 DPBs is then obtained with the standard min-fill-ordering heuritic [18], giving a width of 6. The EteRNA77 puzzle is 105 nts long, and consists in a collection of helices interspersed with destabilizing internal loops. RNApond failed to produce a solution, and its final set of DBPs consists of 183 pairs, for which the same heuristic yields a tree decomposition of width 9. We further make both tree decompositions binary through bag duplications (see Supplementary Section A), giving an FPT runtime to our algorithm, while potentially lowering the number of lost edges.

Executing the tree-diet algorithm (Theorem 7) on both graphs and their tree decompositions, we obtained simplified graphs, having lower treewidth while typically losing few edges, as illustrated and reported in Figure 9. Remarkably, the treewidth of the DBPs inferred for EteRNA22 can be decreased to *tw*′ = 5 by only removing 5 DBPs/edges (460/465 retained), and to *tw*′ = 4 by removing 4 further DBPs (456/465). For EteRNA77, our algorithm reduces the treewidth from 9 to 6 by only removing 7 DBPs.

**Figure 9.**
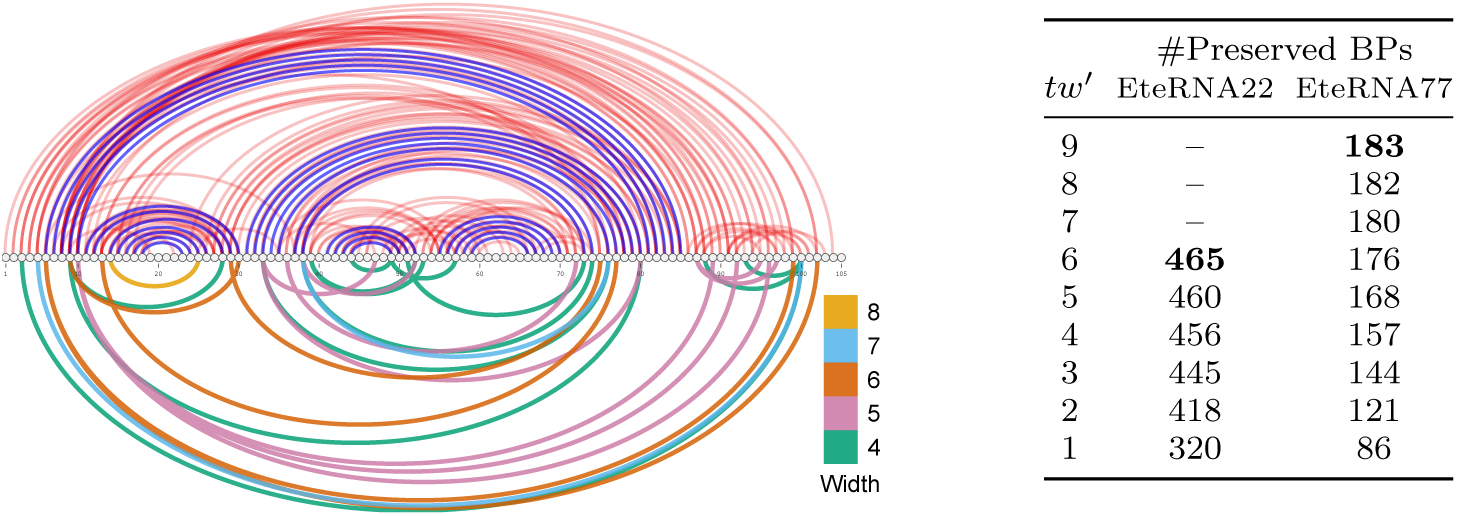
(Left) Target secondary structure (blue BPs), full set of disruptive base pairs (DPB; top) inferred by RNAPond on the Eterna77 puzzle, and subsets of DBPs (bottom) cumulatively removed by the tree-diet algorithm to reach prescribed treewidths. (Right) Number of BPs retained by our algorithm, targeting various treewidth values for the EteRNA22 and EteRNA77 puzzles.

Rough estimates can be provided for the tradeoff between the rejection and preprocessing complexities, by assuming that removing a DBP homogeneously increases the value of 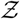 by a factor *α* := 16*/*10 (#pairs/#incomp. pairs). The relative increase in partition function is then 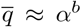, when *b* base pairs are removed. For EteRNA22, reducing the treewidth by 2 units (6*→*4), *i.e*. a 16 fold reduction of the memory and preprocessing time, can be achieved by removing 9 DBPs, *i.e*. a 69 fold expected increase in the time of the generation phase. For EteRNA77, the same 16 fold (*tw*′ = 9 *→* 7) reduction of the preprocessing time/space can be achieved through an estimated 4 fold increase of the generation time. A more aggressive 256 fold memory gain can be achieved at the expense of an estimated 1 152 fold increase in generation time. Given the large typical asymmetry in runtimes and implementation constants between the computation-heavy preprocessing and, relatively light, generation phases, the availability of an algorithm for the tree-diet problem provides new options, especially to circumvent memory limitations.

### 5.2 Structural alignment of complex RNAs

Structural homology is often posited within functional families of non-coding RNAs, and is foundational to algorithmic methods for multiple RNA alignments [13], considering RNA base pairs while aligning distant homologs. In the presence of complex structural features (pseudoknots, base triplets), the sequence-structure alignment problem becomes hard, yet admits XP solutions based on the treewidth of the base pair + backbone graph. In particular, Rinaudo *et al*. [12] describe a Θ(*n.m*^*tw*+1^) algorithm for optimally aligning a structured RNA of length *n* onto a genomic region of length *m*. It optimizes an alignment score that includes: i) substitution costs for matches/mismatches of individual nucleotides and base pairs (including arc-breaking) based on the RIBOSUM matrices [45]; and ii) an affine gap cost model [46]. We used the implementation of the Rinaudo *et al*. algorithm, implemented in the LicoRNA software package [47, 48].

#### 5.2.1 Impact of treewidth on the structural alignment of a riboswitch

In this case study, we used our tree-diet algorithm to modulate the treewidth of complex RNA structures, and investigate the effect of the simplification on the quality and runtimes of structure-sequence alignments. We considered the Cyclic di-GMP-II riboswitch, a regulatory motif found in bacteria that is involved in signal transduction, and undergoes conformational change upon binding the second messenger c-di-GMP-II [49, 50]. A 2.5Å resolution 3D model of the c-di-GMP-II riboswitch in *C. acetobutylicum*, proposed by Smith *et al*. [51] based on X-ray crystallography, was retrieved from the PDB [24] (PDBID: 3Q3Z). We annotated its base pairs geometrically using the DSSR method [52]. The canonical base pairs, supplemented with the backbone connections, were then accumulated in a graph, for which we heuristically computed an initial tree decomposition *𝒯*_4_, having treewidth *tw* = 4.

We simplified our the initial tree decomposition *𝒯*_4_, and obtained simplified models *𝒯*_3_, and *𝒯*_2_, having width *tw*′ = 3 and 2 respectively. As controls, we included tree decompositions based on the secondary structure (max. non-crossing set of BPs; *𝒯*_2*D*_) and sequence (*𝒯*_1*D*_). We used LicoRNA to predict an alignment *a*_*T*, *w*_ of each original/simplified tree decomposition *𝒯* onto each sequence *w* of the c-di-GMP-II riboswitch family in the RFAM database [13] (RF01786). Finally, we reported the LicoRNA runtime, and computed the Sum of Pairs Score (SPS) [53] as a measure of the accuracy of *a*_*T*, *w*_ against a reference alignment 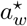 :

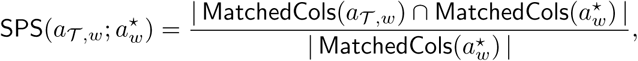

using as reference the alignment 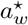 between the 3Q3Z sequence and *w* induced by the manually-curated RFAM alignment of the RF01786 family.

The results, presented in Figure 10, show a limited impact of the simplification on the quality of the predicted alignment, as measured by the SPS in comparison with the RFAM alignment. The best average SPS (77.3%) is achieved by the initial model, having treewidth of 4, but the average difference with simplified models appears very limited (*e.g*. 76.5% for *𝒯*_3_), especially when considering the median. Meanwhile, the runtimes mainly depend on the treewidth, ranging from 1h for *𝒯*_4_ to 300ms for *𝒯*_1*D*_. Overall, *𝒯*_2*D*_ seems to represent the best compromise between runtime and SPS, although its SPS may be artificially inflated by our election of RF01786 as our reference (built from a covariance model, *i.e*. essentially a 2D structure). Finally, the difference in number of edges (and induced SPS) between *𝒯*_2*D*_ and *𝒯*_2_, both having *tw* = 2, exemplifies the difference between the Tree-Diet and Graph-Diet problems, and motivates further work on the latter.

**Figure 10.**
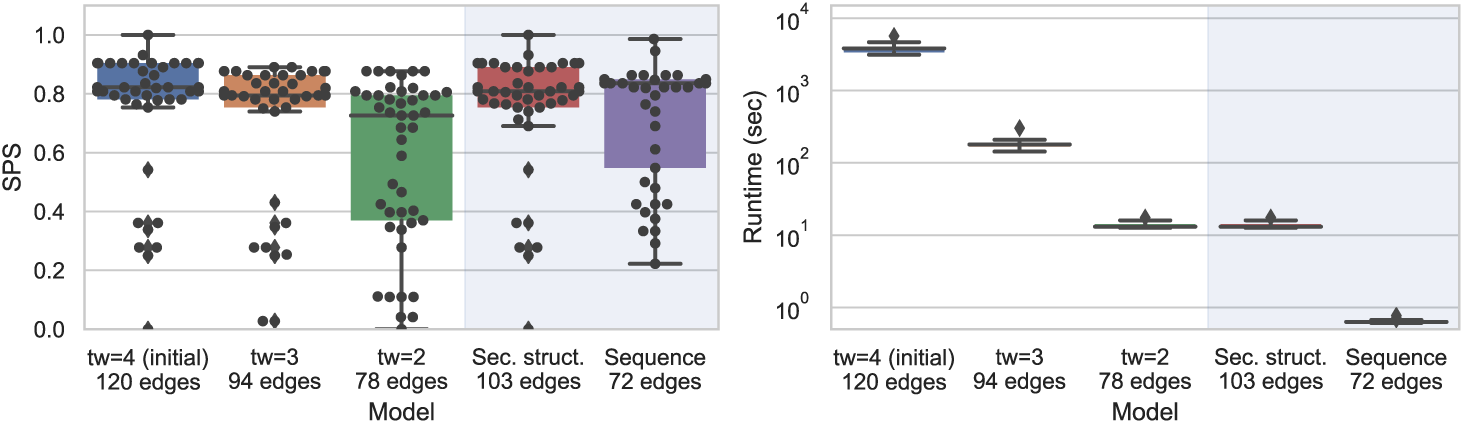
Impact on alignment quality (SPS; Left) and runtime (Right) of simplified instances for the RNA sequence-structure alignment of the pseudoknotted c-di-GMP-II riboswitch. The impact of simplifications on the quality of predicted alignments, using RFAM RF01786 as a reference, appears limited while the runtime improvement is substantial.

#### 5.2.2 Exact iterative strategy for the genomic search of ncRNAs

In this final case study, we consider an exact filtering strategy to search new occurrences of a structured RNA within a given genomic context. In this setting, one attempts to find all *ε*-admissible (cost *≤ ε*) occurrences/hits of a structured RNA *S* of length *n* within a given genome of length *g ≫ n*, broken down in windows of length *κ.n, κ >* 1. Classically, one would align *S* against individual windows, and report those associated with an *ϵ*−admissible alignment cost. This strategy would have an overall Θ(*g · n*^*tw*+2^) time complexity, applying for instance the algorithm of [12].

Our instance simplification framework enables an alternative strategy, that incrementally filters out unsuitable windows based on models of increasing granularity. Indeed, for any given target sequence, the min alignment cost *c*_*δ*_ obtained for a simplified instance of treewidth *tw* − *δ* can be corrected (*cf* Supplementary Section D) into a lower bound 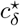 for the min alignment cost 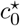 of the full-treewidth instance *tw*. Any window such that 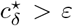 thus also obeys 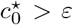, and can be safely discarded from the list of putative *ε*-admissible windows, without having to perform a full-treewidth alignment. Given the exponential growth of the alignment runtime for increasing treewidth values (see Figure 10-right) this strategy is expected to yield substantial runtime savings.

We used this strategy to search occurrences of the Twister ribozyme (PDBID 4OJI), a highly-structured (*tw* = 5) 54nts RNA initially found in *O. sativa* (Asian rice) [54]. We targeted the *S. bicolor* genome (sorghum), focusing on a 10kb region centered on the 2,485,140 position of the 5th chromosome, where an instance of the ribozyme was suspected within an uncharacterized transcript (LOC110435504). The 4OJI sequence and structure were extracted from the 3D model as above, and included into a tree decomposition *𝒯*_5_ (73 edges), simplified into *𝒯*_4_ (71 edges), *𝒯*_3_ (68 edges) and *𝒯*_2_ (61 edges) using the tree-diet algorithm.

We aligned all tree decompositions against all windows of size 58nts using a 13nts offset, and measured the score and runtime of the iterative filtering strategy using a cost cutoff *ε* = −5. The search recovers the suspected occurrence of twister as its best result (Figure 11.C), but produced hits (*cf* Figure 11.D) with comparable sequence conservation that could be the object of further studies. Regarding the filtering strategy, while *𝒯*_2_ only allows to rule out 3 windows out of 769, *𝒯*_3_ allows to eliminate an important proportion of putative targets, retaining only 109 windows, further reduced to 15 windows by *𝒯*_4_, 6 of which end up as final hits for the full model *𝒯*_5_ (*cf* Figure 11.A). The search remains exact, but greatly reduces the overall runtime from 24 hours to 34 minutes (42 fold!).

**Figure 11.**
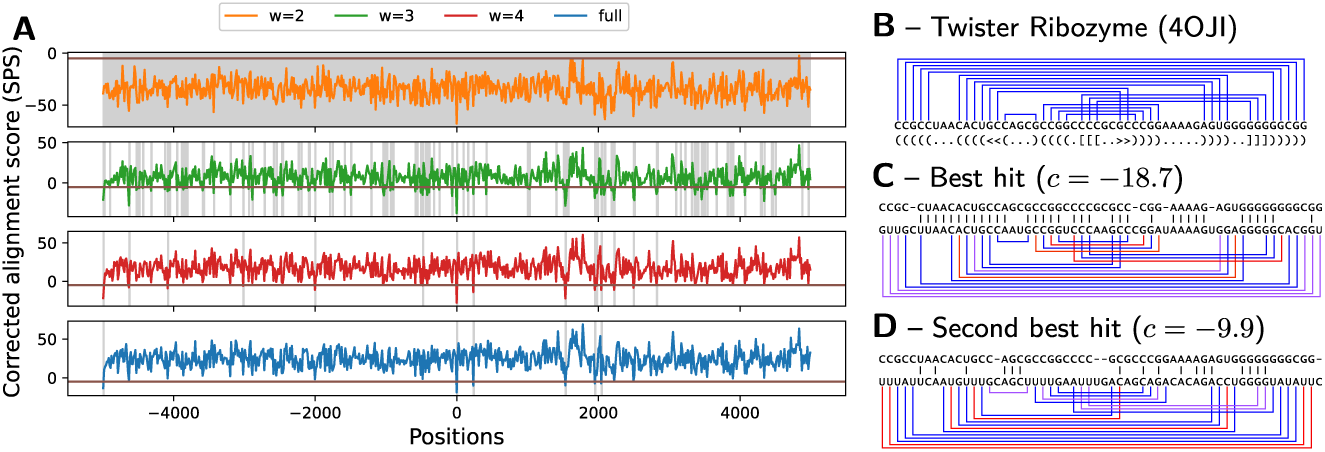
Corrected costs associated with the search for structured homologs of the Twister ribozyme in chromosome 5 of *S. bicolor*, using simplified instances of various treewidth (A). Gray areas represent scores which, upon correction, remain below the cutoff, and have to be considered for further steps of the iterated filtering. Canonical base pairs of the ribozyme (PDBID 4OJI; B), mapped onto to the best hit (C) and second best hit (D) found along the search colored depending on their support in the target sequence (Red: incompatible; Purple: unstable G-U; Blue: stable).

## 6 Conclusion and discussion

We have established the parameterized complexity of three treewidth reduction problems, motivated by applications in Bioinformatics, as well as proposed practical algorithms for instances of reasonable treewidths. The reduced widths obtained by our proposed algorithm can be used to obtain: i) sensitive heuristics, owing to the consideration of a maximal amount of edges/information in the thinned graphs; ii) *a posteriori* approximation ratios, by comparing the potential contribution of removed edges to the optimal score obtained of the thinned instance by a downstream FPT/XP algorithm; iii) substantial practical speedups without loss of correctness, *e.g*. when partial filtering can be safely achieved based on simplified input graphs.

### 6.1 Open questions

Regarding the parameterized complexity of Graph-Diet and Tree-Diet, some questions remain open (see Table 1): an FPT algorithm for Tree-Diet(ideally, with 2^*O*(*tw*)^ *· n* running time), would be the most desirable, if possible satisfying the backbone constraints. The existence of such an algorithm is not trivial. In particular, it is perhaps worth noting that it is not implied by the existence of an FPT algorithm for graph-diet with the input treewidth as a parameter (1). Indeed, in comparison to the latter, tree-diet subtly restricts the search space to tree decompositions that subsets of the input. It follows that the result of graph diet for a graph *G* may substantially differ from the result of tree-diet given a tree decomposition *𝒯* of *G* as input. We also aim at trying to give efficient exact algorithms for graph diet in the context of RNA (we conjecture this is impossible in the general case). Finally, we did not include the number of deleted edges in our multivariate analysis: even though in practice it is more difficult *a priori* to guarantee their small number, we expect it can be used to improve the running time in many cases.

### 6.2 Backbone Preservation

In two of our applications, the RNA secondary structure graph contains two types of edges: those representing the *backbone* of the sequence (i.e., between consecutive bases) and those representing base pair bonds. In practice, we want all backbone edges to be visible in the resulting tree-decomposition, and only base pairs may be lost. This can be integrated to the Tree-Diet model (and to our algorithms) using weighted edges, using the total weight rather than the count of deleted edges for the objective function. Note that some instances might be unrealizable (with no tree-diet preserving the backbone, especially for low *tw*′). In most cases, ad-hoc bag duplications can help avoid this issue. The design of pre-processing methods, involving bag duplications or other operations on tree decompositions, and aimed at ensuring the existence of a backbone-preserving tree-diet will be the subject of future work.

From a theoretical perspective, weighted edges may only increase the algorithmic complexity of the problems. However, a more precise model could consider graphs which already include a hamiltonian path (the backbone), and the remaining edges form a degree-one or two subgraph. Such extra properties may, in some cases, actually reduce the complexity of the problem. As an extreme case, we conjecture the Path-Diet problem for *tw*′ = 1 becomes polynomial in this setting.

## Acknowledgements

The authors would like to thank Julien Baste for pointing out prior work on treewidth modulators, and providing valuable input regarding vertex deletion problems.

## Availability of data and materials

Source code of tree-diet method available at:https://gitlab.inria.fr/amibio/tree-diet

## Supplementary Material

### Section A: Editing Trees before the Diet

Any tree decomposition can be transformed into a binary one through the duplications of bags having more than 2 children. To do so in practice, one will, as long as the tree decomposition is not binary, apply the following transformation:

1. Find a bag *X* with children *Y*_1_, …, *Y*_Δ_ and Δ *>* 2.
2. Introduce a new bag *X*′ with the same content as *X* and locally modify the tree decomposition in the following way: *X* will now have *Y*_1_ and *X*′ as children, while *X*′ will have *Y*_2_ *· · · Y*_Δ_.

When it is no longer possible to apply this transformation, the tree decomposition is binary. For each bag having originally Δ *>* 2 children in the decomposition, Δ − 1 new bags have been introduced. In total, with *N*_*bags*_ the original number of bags in the decomposition, strictly less than *N*_*bags*_ new bags have been introduced (each new bag is associated to an edge of the original tree decomposition).

This tranformation is in fact the first step towards obtaining a *nice tree decomposition* [42, 29].

A question that arises then is what impact these modifications may have on the output of Tree-Diet, when applied to the tree decomposition given as input. We argue that duplication operations (as used above to get a binary tree decomposition) can only improve the solution, i.e decrease the number of *lost* edges. Indeed, within the coloring formulation of the problem, new bags yield new opportunities for an edge to be *represented*, with both its end-points green in some bag. See Figure 12 for an illustration.

**Figure 12.**
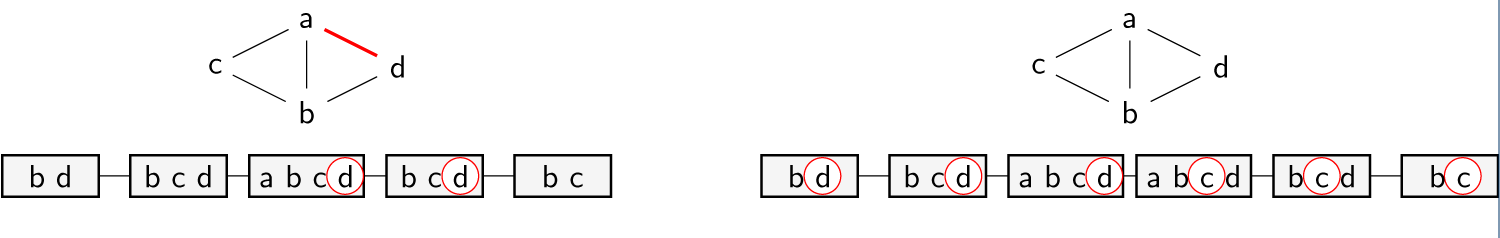
Left: A graph and a path-decomposition whose optimal 1-tree-diet loses an edge (ad). However, duplicating the bag abcd (right) yields a tree-decomposition with a lossless 1-tree-diet.

More generally, any operation on the input tree decomposition that does not suppress any of the original bags can only improve the solution to the Tree Diet problem. We do not tackle here the problem of finding the best edition operations to apply onto a tree decomposition given as input to Tree Diet, which is an a priori difficult task.

### Section B: Pseudo-code

Algorithm 1 and 2 present a pseudo-code of our dynamic programming algorithm for Tree Diet, with a memoization approach. The C++/pybind11 [44] implementation is available at https://gitlab.inria.fr/amibio/tree-diet.

Note that the implementation allows to solve a more general *weighted* version of Tree Diet, where each edge is given a weight, and the objective is to find a (*tw*−*tw*′)-diet of the input tree decomposition preserving a set of edges of maximum total weight.

In the context of RNA applications, this feature allows to favour as much as possible preservation of the backbone of RNA molecules, i.e. edges between consecutive nucleotides along the string, by assigning them a weight greater than the number of non-backbone edges.

Edge weights are passed to the function in the form of a dictionary/map *W* associating a real weight to each edge. Within Algorithm 1, the only place where it is taken into account is the the *count* function, which computes the weight of edges accounted for by the bag that is currently visited.

#### Algorithm 1: Dynamic programming algorithm for Tree-Diet.

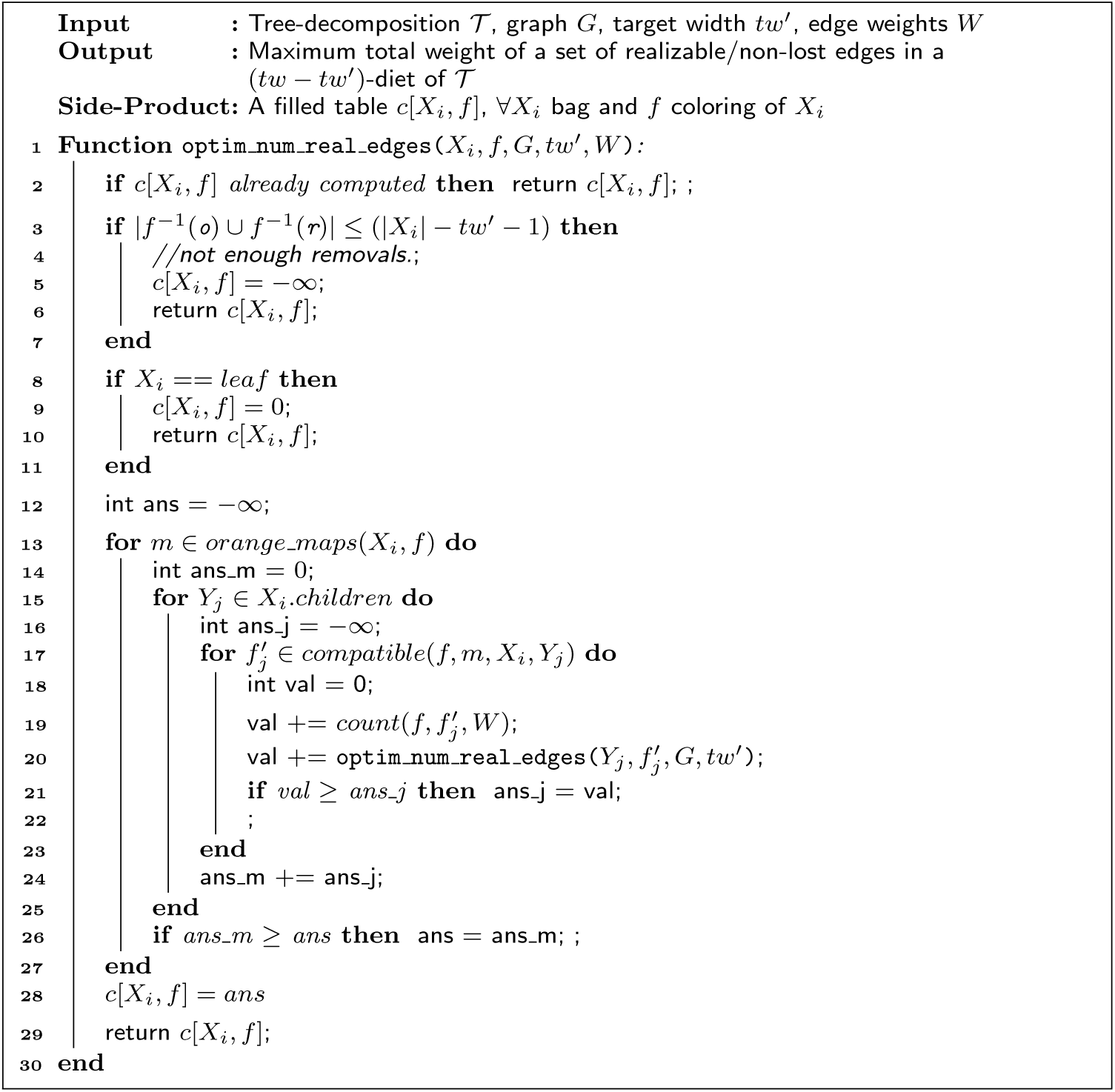

### Section C: Correctness of the rejection-based sampling of RNA designs

A recent method for RNA design, called RNAPond [22], implements a sampling approach to tackle the inverse folding of RNA. Targeting a secondary structure *S* of length *n*, it performs a Boltzmann-weighted sampling of sequences and, at each iteration, identifies Disruptive Base Pairs (DBPs) that are not in *S*, yet are recurrent in the Boltzmann ensemble of generated sequences. Those base pairs are then added to a set *𝒟* of DBPs, and excluded in subsequent generations through an assignment of non-binding pairs of nucleotides, outside of *B* := *{*(G, C), (C, G), (A, U), (U, A), (G, U), (U, G)*}*.

#### Algorithm 2: Backtracking procedure for Tree-Diet.

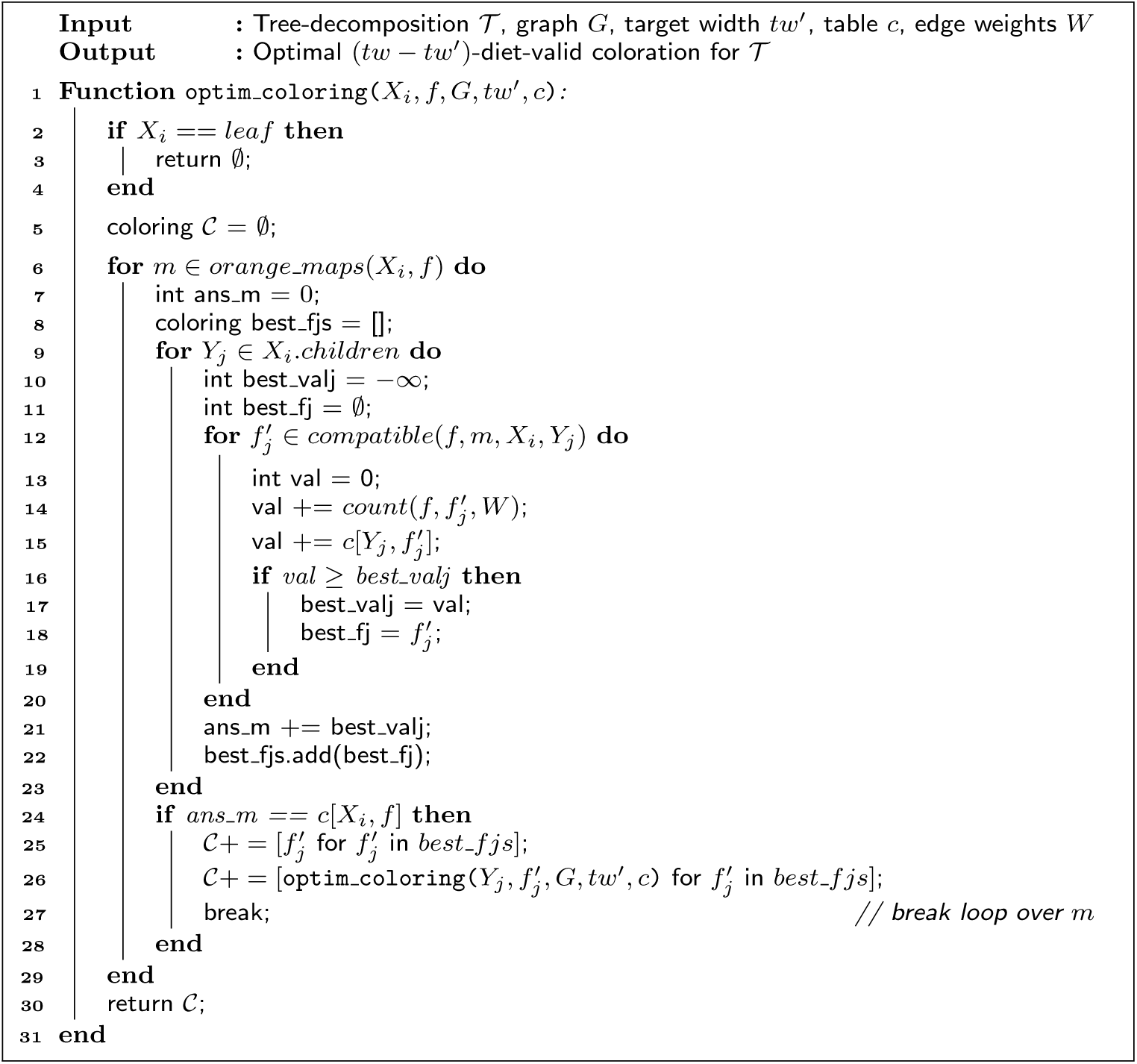

At the core of the method, one finds a random generation algorithm which takes as input a secondary structure *S* and a set *𝒟* of DBPs. The algorithm generates from the set *W*_*S,D*_ of sequences *w ∈ {*A, C, G, U*}*^*n*^ which are: i) compatible with all (*i, j*) *∈ S, i.e*. (*w*_*i*_, *w*_*j*_) *∈ ℬ*; and ii) incompatible with all (*k, l*) *∈ 𝒟, i.e*. (*w*_*k*_, *w*_*l*_) *∉ ℬ*. The algorithm then enforces a (dual) Boltzmann distribution over the sequences in *𝒲*_*S,𝒟*_:

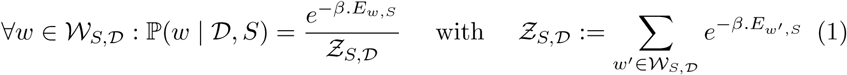

where *β >* 0 is an arbitrary constant akin to a temperature. Yao *et al*. describe an algorithm which generates *k* sequences in Θ(*k*(*n* + |*𝒟*|)) time, after a preprocessing in Θ(*n*.|*𝒟*|.4^*tw*^) time and Θ(*n*.4^*tw*^) space, where *tw* is the treewidth of the graph having edges in *S ∪ 𝒟*.

The discrepancy in the preprocessing and sampling complexities suggests an alternative strategy, utilizing rejection on top of a relaxed sampling. Namely, we consider a rejection algorithm, which starts from a relaxation (*S*′, *𝒟*′) of the initial constraints (*S*′ *∪ 𝒟*′ *⊂ S ∪ 𝒟*), and iterates Yao *et al*.’s algorithm to generate sequences in *W*_*S ′*_,_*𝒟 ′*_*⊃ 𝒲*_*S,𝒟*_, rejecting those outside of *𝒲*_*S,𝒟*_, until *k* suitable ones are obtained. The rejection algorithm generates a given sequence *w ∈ 𝒲*_*S,𝒟*_ on its first attempt with probability 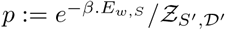 and, more generally, after *r* rejections with probability (1 − *q*)^*r*^ *p* with 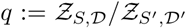. The overall probability of emitting *w* is thus

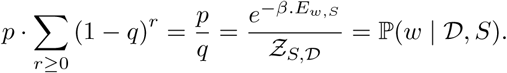

In other words, our relaxed generator coupled with the rejection step, represents an unbiased algorithm for the Boltzmann distribution of Eq. (1) over *𝒲*_*S*, *D*_.

Meanwhile, the average-case complexity can be impacted by the strategy. Indeed, the relaxed instance (*S*′, *𝒟*′) can accelerate the preprocessing due to a reduced treewidth *tw*′ *≤ tw*. The rejection step only increases the expected number of generations by a factor 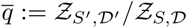, representing the inflation of the sequence space, induced by the relaxation of the constraints. Overall, the average-case time complexity of the rejection algorithm is in 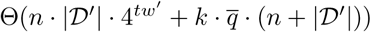 time and Θ(*n ·* 4^*tw ′*^) space. This space improvement is notable when *tw*′ *< tw*, and could be key for the practical applicability of the method, especially given that memory represents the bottleneck of most treewidth-based DP algorithms.

### Section D: Lower bound for the min. alignment cost from simplified models

Here, we justify the filtering strategy described in Section 5.2.2. Namely, we formally prove that, given a structured RNA *S* and a targeted genomic region *w*, a lower bound for the minimal alignment cost of *S* and *w* can be obtained from the minimal alignment cost of some *S*′ *⊆ S* and *w*. If this lower bound for *S*′ *⊆ S* is higher than the specified cutoff *ε*, then there is no need to align *w* to the full model *S*, as the resulting cost is guaranteed to stay above the selection cutoff *ε*.

Let *S* be an arc-annotated sequence of length *m* (*S*_*i*_ denotes the *i*th character of *S*), *w* be a target (flat) sequence of length *m*, and *μ* : [1, *n*] *→* [1, *m*] *∪ {⊥}* represents an alignment^[1]^. We consider the following cost function, adapted from eciteRinaudo2012, which quantifies the quality of an alignment *μ* for *S* and *w*:

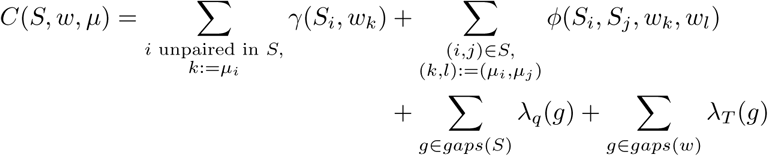

where

- *γ*(*a, b*) returns the *substitution cost* which penalizes (mismatches) or rewards (matches) the substitution of *a* into *b* (set to 0 and handled in gaps if *b* =⊥);
- *ϕ*(*a, b, c, d*) returns a *base pair substitution cost*, penalizing (arc breaking) or rewarding (conservation or compensatory mutations) the transformation of nucleotides (*a, b*) into nucleotides/gaps (*c, d*) (set to 0 and handled in gaps if (*c, d*) = (⊥, ⊥));
- *λ*_*S*_ and *λ*_*T*_ penalize gaps introduced by *μ* respectively in *S* and *w* (affine cost model).

Given this definition, consider a simplified model *S*′ *⊂ S*, associated with a minimal cost

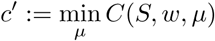

and denote by *c*^*^ the minimal cost of the full model *S*, we have the following inequality.

#### Proposition 3

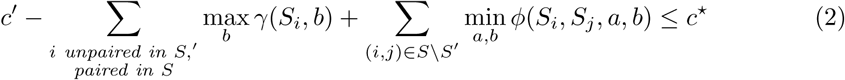

*Proof* For any alignment, we have, per the definition of *C*(*S, w, μ*):

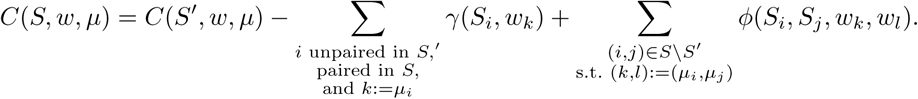

Minimizing over all alignment *μ*, one obtains

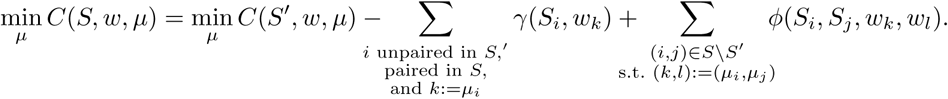

Independently minimizing each term of the right-hand-side, we obtain a first lower bound

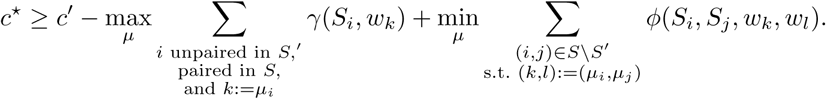

further coarsened by an independent optimization of the elements in the sums

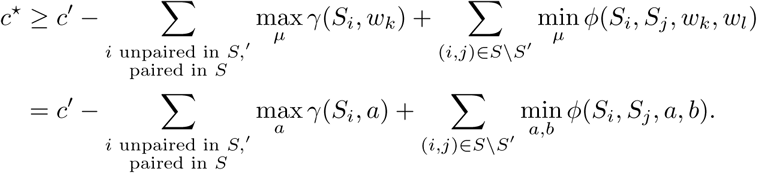

where the last line is obtained by considering the worst-case contributors to nucleotides and base pairs substitutions. Importantly, the right-hand side no longer depends on *μ* any more, and can be used to easily computed a corrected score/lower bound. □

The corrected expression, shown in the left hand side of Equation (2) allows, when lower than a cutoff *ε*, to safely discard *w* as a potential hit for the full model *S*. This corrected score is plotted in Figure 11A, allowing for a gradual reduction of the search space for *ε*-admissible hits. We show in Figure 13 the corrected scores obtained for simplified structures *S*′ of various treewidths, plotted against the scores of the full target structure.

**Figure 13.**
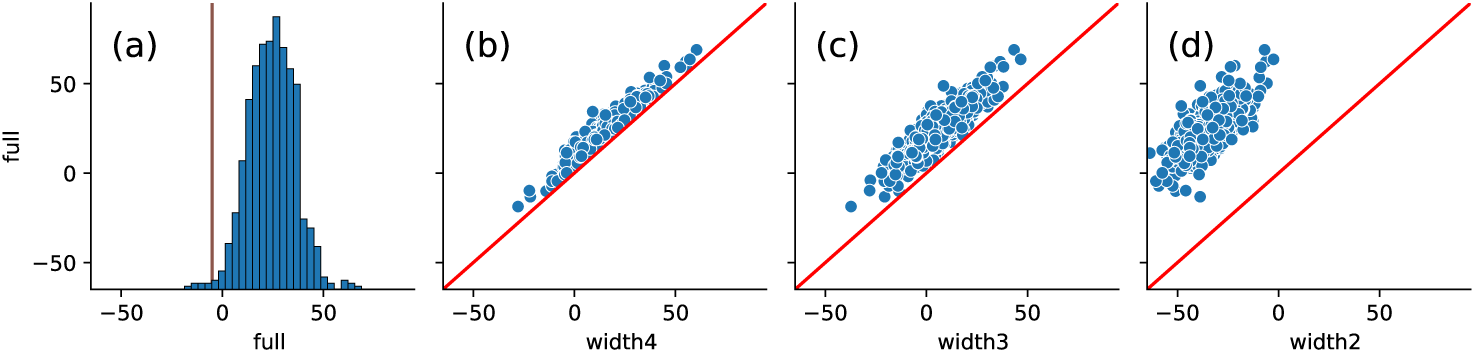
(a) Histogram of alignment scores obtained by aligning the full structure (*tw* = 5) model of the Twister ribozyme (pdb-id: 4OJI) with *κ · n*-sized windows in a 10kb region of the 5^*th*^ chromosome of *S. bicolor*. A vertical line is positioned at the *ϵ* threshold. (b;c;d) Corrected alignment scores obtained for reduced-treewidth models for each window, plotted against the corresponding score of the full model. The corrected alignment score indeed acts as a lower bound to the full-model score (points above the *y* = *x* red line), allowing an iterative filtering strategy.

## Footnotes

[1] An alignment _*μ*_ is subject to further constraints, notably including some restricted form of monotonicity, when represented as a function. However, those constraints are reasonably intuitive and we omit them in this discussion for the sake of simplicity.

